# Intrinsic Circadian Organization of the Spinal Cord and Dorsal Root Ganglia

**DOI:** 10.64898/2026.01.27.702043

**Authors:** Shintaro Yamazaki, Lee Wulund, Utham K. Valekunja, Akhilesh B. Reddy

**Affiliations:** Department of Systems Pharmacology & Translational Therapeutics, Perelman School of Medicine, University of Pennsylvania, Philadelphia, PA 19104, USA; Institute for Translational Medicine and Therapeutics, Perelman School of Medicine, University of Pennsylvania, Philadelphia, PA 19104, USA; Chronobiology and Sleep institute (CSI), Perelman School of Medicine, University of Pennsylvania, Philadelphia, PA 19104, USA; University of Cambridge Metabolic Research Laboratories, Wellcome Trust-MRC Institute of Metabolic Science, Addenbrooke’s Hospital, Cambridge CB2 0QQ, UK

## Abstract

Circadian rhythms coordinate daily fluctuations in physiology and behavior, yet their organization within primary sensory pathways remains poorly defined. Although somatosensory responsiveness varies across the day-night cycle, it is unclear whether peripheral sensory circuits possess molecular mechanisms for temporal regulation. Here, we demonstrate that the spinal-peripheral sensory axis harbors robust, tissue-autonomous circadian clocks. Using real-time bioluminescence imaging, we observed sustained oscillations of the core clock protein PER2 in the spinal dorsal horn and dorsal root ganglia (DRGs), indicating autonomous circadian timing within these tissues. To define the molecular scope of this regulation, we performed RNA sequencing across a 52-hour circadian time course in DRGs. Circadian analysis identified 626 rhythmic transcripts, representing 3.6% of expressed genes. These genes exhibited non-uniform phase distributions and segregated into discrete temporal clusters. Functional annotation revealed phase-specific enrichment of biological processes related to transport, neuronal structure, and proteostasis, suggesting coordinated temporal deployment of distinct molecular programs rather than uniform oscillations across the circadian cycle. Cross-referencing circadian genes with neuropathic pain–associated gene sets revealed limited overlap; however, overlapping genes aligned to specific baseline phase windows enriched for regenerative annotations. Potassium channel-related signaling components implicated in neuropathic pain also showed baseline circadian modulation. Together, these findings establish the spinal dorsal horn and DRGs as intrinsically circadian tissues and reveal a temporally structured molecular landscape in primary sensory neurons, providing a framework for understanding how peripheral sensory processing, plasticity, and homeostatic regulation are coordinated across the day-night cycle.

## Introduction

Circadian rhythms are endogenous oscillations that synchronize molecular, cellular, and physiological processes to the 24-hour light-dark cycle. These rhythms are generated by intracellular timekeeping mechanisms composed of core clock genes arranged in transcription-translation negative feedback loops, producing near-24-hour oscillations in gene expression and physiology ^1^. Individual cells in the mammalian brain and peripheral tissues are capable of autonomous circadian oscillations through rhythmic expression of core clock genes ^2^, while coordination across tissues is achieved via systemic cues originating from the master circadian pacemaker located in the suprachiasmatic nucleus (SCN) of the hypothalamus. Accordingly, circadian regulation is now recognized as a fundamental organizing principle across diverse organ systems ^1^.

The primary somatosensory system lies at the interface between the external environment and the central nervous system (CNS). Peripheral sensory information is conveyed by primary afferent neurons whose cell bodies reside in the dorsal root ganglia (DRGs), with bifurcating axons projecting to peripheral targets and centrally to the dorsal horn of the spinal cord ^3^. DRG neurons comprise molecularly distinct subtypes specialized for mechanosensation, thermosensation, proprioception, and nociception, exhibiting diverse excitability profiles and ion channel expression ^4–7^. Although lacking local synaptic integration, DRGs are embedded in a specialized glial, immune, and vascular microenvironment that modulates neuronal excitability and plasticity, positioning the ganglion as an active regulator rather than a passive relay of sensory information ^8^.

Temporal variation in somatosensory responsiveness has been recognized for decades. Human studies dating back to the 1970s reported diurnal fluctuations in pain perception across clinical contexts, as well as time-of-day differences in analgesic efficacy ^9–14^. Similarly, rodent studies demonstrate circadian modulation of thermal and mechanical sensitivity, and in chemotherapy-induced disease models, although the phase and magnitude of these rhythms vary across experimental paradigms ^15–21^. Notably, rhythmic variation in sensory thresholds persists in models of nerve injury ^22^, suggesting that time-of-day effects are not solely driven by behavioral state or environmental cues. At the molecular level, components of nociceptive signaling pathways also exhibit 24-hour variation. For example, rhythmic expression of *Tac1*, encoding the neurotransmitter Substance P, oscillates in the mouse spinal cord and dorsal horn^23^. Diurnal oscillations of the α2δ-1 subunit of voltage-dependent calcium channels, as well as time-of-day variation in gabapentin binding capacity, have also been reported in DRGs following partial sciatic nerve ligation ^20^.

Despite this body of work, a critical gap remains: whether temporal variation in somatosensory processing arises from intrinsic circadian organization within the primary sensory pathways themselves. In particular, the existence, robustness, and molecular mechanisms of autonomous circadian clocks within the spinal cord and DRGs have not been systematically characterized. Here, we investigate circadian clock function within the mouse spinal cord dorsal horn and DRGs using real-time bioluminescence imaging and transcriptomic profiling. Using *mPer2^luc^* reporter mice, we assess the presence and stability of intrinsic circadian oscillations in sensory tissues ^24^. We further define the molecular landscape of circadian regulation in DRGs using RNA sequencing across a circadian time course. Together, these approaches demonstrate that peripheral sensory pathways harbor robust intrinsic circadian clocks and exhibit widespread temporal organization of gene expression. These findings establish circadian regulation as a fundamental molecular feature of the spinal-peripheral sensory axis and provide a framework for understanding how time-of-day information shapes somatosensory organization, with implications for the optimization of pain-related interventions.

## Methods

### Animals

All animal procedures were conducted in accordance with the UK Animals (Scientific Procedures) Act 1986, and approved by the UK Home Office and the local ethical review panel at the University of Cambridge. Adult and postnatal female and male C57Bl/6J and *mPer2^Luc^*^24^ reporter mice, were used in this study. Unless otherwise stated, mice were bred and maintained under controlled environmental conditions with *ad libitum* access to food and water, and housed on a 12 h:12 h light-dark (LD) cycle. Ambient temperature was monitored daily and maintained at 22 ± 2 °C.

### Organotypic tissue culture and bioluminescence recordings

Organotypic tissue cultures were prepared from adult and postnatal (P7-P10) *mPer2^Luc^* reporter mice. Mice were humanely culled by cervical dislocation, and tissues were rapidly dissected into cold dissection medium consisting of Gey’s Balanced Salt Solution (Sigma-Aldrich, St Louis, MO, USA) supplemented with 5.0 g/L glucose (Sigma-Aldrich), 100 nM MK-801 (Sigma-Aldrich), 50 μM DL-AP5 (Tocris Bioscience, Bristol, UK), and 3 mM MgCl2 (Sigma-Aldrich).

Spinal cords were isolated and sectioned in the transverse plane at 300 µm thickness using a McIlwain tissue chopper (Mickle Laboratory Engineering, Guildford, UK). SCN slices were prepared in the coronal plane at 200 µm thickness using a vibrating microtome (7000smz-2, Campden Instruments, Loughborough, UK). For comparisons across spinal levels, spinal cords were bisected longitudinally along the midline and cultured intact. Adult mouse DRGs were dissected using fine forceps, with both proximal and distal axons severed.

All tissue sections were placed on culture membranes (Merck Millipore, Watford, UK) and maintained in sealed culture dishes containing modified air medium composed of DMEM (Sigma-Aldrich) supplemented with 30 mM glucose, 20 mM HEPES (Sigma-Aldrich), 0.035% NaHCO_3_ (Sigma-Aldrich), 25% (v/v) heat-inactivated horse serum (Thermo Fisher Scientific, Waltham, MA, USA), 1× GlutaMAX™ (Thermo Fisher Scientific), 1× B-27® supplement (Thermo Fisher Scientific), 1 mM luciferin (Biosynth, Staad, Switzerland), and 1× MycoZap™ Plus-PR (Lonza, Basel, Switzerland).

Bioluminescence recordings were performed in a custom-made imaging system (Cairn Research Ltd, Faversham, UK) equipped with an Andor iKon-M 934 CCD camera (Andor Technology, Belfast, UK) housed within a Galaxy 170R incubator (Eppendorf, Hamburg, Germany). Temperature control and image acquisition were managed using MetaMorph Software (Molecular Devices, Sunnyvale, CA, USA). Images were acquired using 25-minute exposure times at every 30-minute intervals. Image stacks were compiled in Image J (NIH, Bethesda, MD, USA), and bioluminescence signals were quantified by manually defined region of interest (ROIs) encompassing each tissue slice. Integrated density values were extracted and analysed using the CellulaRhythm R script ^25^.

High magnification (10×) bioluminescence imaging of spinal cord sections was performed using a Hamamatsu ImageEM 1K camera (Hamamatsu Photonics, Hamamatsu, Japan) mounted on an Eclipse Ti-E microscope (Nikon, Tokyo, Japan). Culture membranes were transferred to 35-mm glass-bottom dishes and maintained in a temperature-controlled incubation chamber (37°C; Okolab, Pozzuoli, Italy). Images were acquired every 30 minutes using NIS Elements AR software (version 4; Nikon), exported to ImageJ, and analyzed as described above.

### Circadian entrainment and RNA isolation

Circadian entrainment was performed in custom-built, ventilated housing cabinets (Tecniplast, UK) with lighting controlled by ClockLab software (Actimetrics, Wilmette, IL, USA). Adult male C57Bl/6J mice were entrained to a 12 h:12 h light-dark (LD) cycle for two weeks, followed by release into constant darkness (DD) for 2 days to assess endogenous circadian rhythms. On the third day in DD, mice were culled by cervical dislocation at 4-h intervals beginning at circadian time 0 (CT0; n=3) and continuing through CT52. CT0 was defined as the projected time of lights-on based on the preceding light-dark cycle.

All DRGs from each animal were rapidly dissected and immediately homogenized in TRI Reagent (Zymo Research, Irvine, CA, USA), snap-frozen on dry ice, and stored at −80°C until processing. For RNA extraction, samples were thawed and transferred to bead-beater tubes (Lysing Matrix D, MP Biomedicals, Santa Ana, CA, USA) and homogenized using a FastPrep-24 instrument (MP Biomedicals; 2 × 30 s at 4.0 m/s). Homogenates were transferred to Phase Lock Heavy Gels (5 Prime, Hilden, Germany), mixed with chloroform (Sigma-Aldrich), and subjected to phenol-chloroform extraction. Following phase separation, ethanol (Sigma-Aldrich) was added to the aqueous phase, and RNA with the Zymo Clean & Concentrator-5 Kit (Zymo Research) according to the manufacturer’s instructions. RNA was eluted in nuclease-free water and stored at −80°C until further processing.

### Quantitative PCR

Total RNA (1 μg per sample) was reverse-transcribed into complementary DNA (cDNA) using the High Capacity cDNA Reverse Transcription Kit (Thermo Fisher Scientific) according to the manufacturer’s instructions. Quantitative PCR (qPCR) was performed using the Universal ProbeLibrary system (Roche, Basel, Switzerland), with primers designed using the Roche online assay design tool. Reactions were run in duplicate using Taqman Gene Expression Master Mix (Thermo Fisher Scientific) on an Applied Biosystems 7900HT Real-Time PCR System (Thermo Fisher Scientific). Relative mRNA expression levels were calculated using the 2^-ΔΔCt^ method and normalized to 18S ribosomal RNA as an internal control. The following primer and probe sets were used: 18S fwd: 5’ gcaattattccccatgaacg, 18S rev: 5’ gggacttaatcaacgcaagc, 18S probe: 48; Per2 fwd: 5’ tccgagtatatcgtgaagaacg, Per2 rev: 5’ caggatcttcccagaaacca, Per2 probe: 5; Bmal1 fwd: 5’ gccccaccgacctactct, Bmal1 rev: 5’ tgtctgtgtccatactttcttgg, Bmal1 probe: 10; Reverba fwd: 5’ cagcatgatcaggtcaatctgt, Reverba rev: 5’ agcaaatcgtaccattaaaacctc, Reverba probe: 95.

### RNA sequencing and analysis

Bulk RNA sequencing was performed on DRG tissue collected across a 52-hour circadian time series (14 time points at 4-hour intervals). For each time point, 2 µg of total RNA was pooled for library preparation using TruSeq® Stranded mRNA Library Prep Kit (Illumina, San Diego, CA, USA). Sequencing was performed on an Illumina HiSeq platform (Illumina) paired-end 50 bp reads.

For analysis, raw sequencing pair-end reads were filtered to remove ribosomal RNA contamination and aligned to the *Mus musculus* reference genome (GRCm39) using TopHat2 (v2.0.12) ^26^, a splice-aware aligner built on Bowtie2. Gene annotations were obtained from Ensembl release 105 (GENCODE M28 equivalent). Alignments were performed independently for each time point, and resulting BAM files were coordinate-sorted using SAMtools (v0.1.19).

Gene- and isoform-level mRNA expression was quantified using the Cufflinks suite (v2.2.1). Expression values were estimated as fragments per kilobase of exon per million mapped reads (FPKM) using Cuffdiff, with bias correction enabled and geometric normalization applied across samples. All time points were processed jointly to generate a unified expression matrix for downstream circadian analysis. Genes with low expression (FPKM ≤ 0.5 across all time points) were excluded from further analysis.

### Circadian rhythmicity analysis

Circadian rhythmicity of expressed genes was assessed using multiple complementary algorithms implemented in R, including RAIN (Rhythmicity Analysis Incorporating Non-parametric methods) ^27^ and Autoregressive Regression based on Spectral Estimation and Regression (ARS/ARSER) ^28, 29^. RAIN was used as the primary method for identifying rhythmic genes due to its robustness to waveform shape and non-sinusoidal oscillations ^27^. P-values were adjusted for multiple testing across all gene expression profiles using the Benjamini-Hochberg procedure, and genes with adjusted RAIN q < 0.15 were considered rhythmic. Phase estimates used for downstream analyses were derived from ARS-adjusted phase values, which provide stable peak-time estimates across noisy time series.

To evaluate whether the observed number of rhythmic genes exceeded that expected by chance, we performed a permutation-based enrichment analysis ^30, 31^. Gene-level circadian rhythmicity was reassessed under a null model generated by randomizing the association between time labels and expression profiles while preserving the overall structure of the dataset. Specifically, for each permutation, expression profiles were randomly reassigned across genes, after which RAIN was reapplied using identical parameters to those used for the observed data. The number of genes passing the rhythmicity threshold (RAIN q < 0.15) was recorded for each permutation. A total of 1,000 permutations were performed to generate an empirical null distribution of rhythmic gene counts. The empirical p-value was calculated as (*K* + 1)/(*N* + 1), where *K* is the number of permutations yielding a rhythmic gene count greater than or equal to the observed value, and *N* is the total number of permutations. This approach provides a conservative estimate of significance and avoids assumptions about the underlying distribution of rhythmicity scores.

### Functional annotation and enrichment analysis

Rhythmic genes were stratified by peak circadian phase and grouped into discrete phase bins for functional analysis. Gene Ontology (GO) classification and enrichment analyses were performed using the clusterProfiler package (version 4.14.6) in R, with *Mus musculus* gene annotations from org.Mm.eg.db (version 3.20.0). Over-representation analysis was conducted using enrichGO with all expressed genes as the background universe. Multiple testing correction was applied using the Benjamini-Hochberg method. Unless otherwise stated, GO terms with adjusted p-values (Benjamini-Hochberg) < 0.1 were considered enriched. For selected analyses requiring stricter stringency (Fig.5F), GO terms were required to meet adjusted p < 0.05 and q < 0.05.

Rhythmic genes were grouped into phase bins based on adjusted ARS phase estimates. To determine an appropriate temporal resolution for phase-resolved analyses, circular variance of gene phases was evaluated across multiple bin widths (2, 3, 4, and 6 h) using circular statistics. Circular variance provides a measure of phase dispersion, with lower values indicating tighter phase coherence. While 2-h bins exhibited the lowest variance, many bins contained insufficient gene numbers for stable downstream functional enrichment. In contrast, 6-h bins showed substantially increased circular variance, consistent with phase mixing of distinct transcriptional waves. Three-hour bins provided an optimal balance, maintaining low circular variance while preserving adequate gene counts for robust gene ontology analysis. All phase-resolved analyses therefore used 3-h bins unless otherwise stated.

## Results

### Spinal dorsal horn exhibits robust intrinsic circadian oscillations

To determine whether primary sensory pathways harbor intrinsic circadian rhythmicity, we examined the mouse cervical spinal cord, a key relay and processing site for ascending somatosensory input. To isolate circadian clock activity from systemic cues, we prepared organotypic spinal cord slices from *mPer2^luc^* reporter mice and maintained them *ex vivo*, where PER2-driven bioluminescence provides a real-time readout of circadian transcriptional activity at the tissue level. Bioluminescence imaging of transverse cervical spinal cord sections from postnatal (P7-P10) mice revealed self-sustained circadian oscillations of *mPer2^luc^* signal within the spinal cord. Rhythmic activity was detected in both dorsal and ventral horn regions; however, oscillation amplitude was consistently higher in the dorsal horn (Figure 1A, B), the primary site of somatosensory afferent input from the periphery. In contrast, ventral horn regions, which predominantly contain motor efferent neurons, exhibited lower-amplitude rhythms.

**Figure 1.**
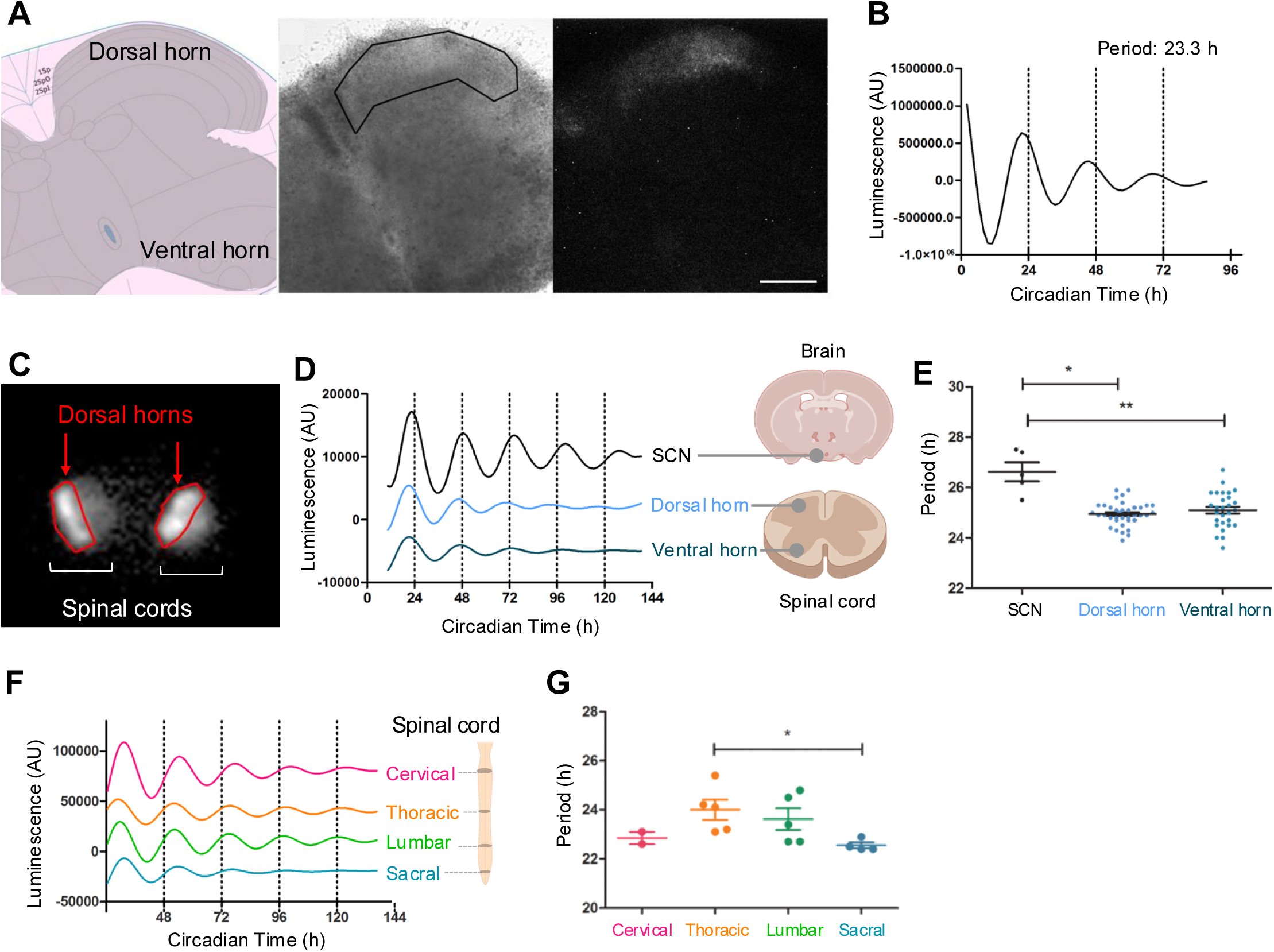
Autonomous PER2 circadian rhythms in the spinal cord dorsal horn. (A) Schematic transverse section of the mouse spinal cord at the C3 level (left), adapted from the Mouse Spinal Cord Atlas, illustrating gray matter (gray) and surrounding white matter (pink), with dorsal and ventral horns indicated. Representative brightfield image (middle) and corresponding bioluminescence image (right) of an organotypic cervical spinal cord slice from mPer2Luc mice. The outlined region marks laminae I–II of the dorsal horn. Scale bar, 200 μm. (B) Representative detrended PER2 bioluminescence trace from the dorsal horn of a cervical spinal cord slice. (C) Macroscopic bioluminescence image of transverse spinal cord slices, with dorsal horn regions outlined in red. (D) Representative detrended PER2 bioluminescence traces from dorsal horn and ventral horn regions of spinal cord slices compared with suprachiasmatic nucleus (SCN) slices from the same animal. (E) Period of PER2 bioluminescence oscillations in SCN, dorsal horn, and ventral horn regions. Data are mean ± SEM with individual data points shown (n = 5 mice; n = 37 dorsal horn slices; n = 28 ventral horn slices). *P < 0.05, **P < 0.01 (Kruskal–Wallis test). (F) Representative detrended PER2 bioluminescence traces from dorsal horn regions of spinal cord slices at cervical, thoracic, lumbar, and sacral levels. (G) Period of PER2 bioluminescence oscillations across spinal levels. Data are mean ± SEM with individual data points shown. *P < 0.05 (Kruskal–Wallis test).

We next compared circadian clock activity in the spinal cord with that of the SCN, which serves as the master circadian pacemaker and exhibits high-amplitude, network-stabilized oscillations *ex vivo*. Using custom-built imaging systems that enabled simultaneous long-term recording from multiple tissue slices, we measured PER2-driven bioluminescence in dorsal horn (n=37 slices), ventral horn (n=28 slices), and SCN (n=5 slices) from the same cohort of animals (n=5 animals) (Figure 1C). Under constant *ex vivo* conditions, spinal cord slices exhibited autonomous circadian oscillations that persisted for multiple cycles, demonstrating intrinsic, tissue-autonomous clock function. Compared to the SCN, oscillations in both the dorsal and ventral horns displayed reduced amplitude and progressive damping over time (Figure 1D), resembling rhythms observed in peripheral organs and extra-SCN brain regions ^32^.

Despite these amplitude differences, spinal cord rhythms maintained stable circadian periodicity (Figure 1E). The intrinsic periods of dorsal horn and ventral horn oscillations were significantly shorter than those of the SCN (24.95 ± 0.08 h and 25.10 ± 0.14 h, respectively, versus 26.62 ± 0.38 h for SCN), consistent with the absence of the specialized intercellular coupling mechanisms that lengthen and stabilize circadian period within the SCN network ^33^.

To assess whether circadian properties varied along the rostrocaudal axis, we the PER2 reporter bioluminescence from dorsal horn slices in the cervical, thoracic, lumbar, and sacral spinal levels. All regions exhibited self-sustained circadian oscillations; however, oscillation amplitudes varied across levels (Figure 1F). In contrast, intrinsic circadian period varied only modestly, with thoracic segments exhibiting slightly longer periods than sacral segments (Figure 1G). These quantitative differences may reflect regional heterogeneity in cellular composition or local circuit properties across spinal levels ^34^, whereas the overall presence of stable circadian rhythms indicates well-maintained intrinsic clock function throughout the spinal dorsal horn.

### Dorsal root ganglia exhibit autonomous circadian clock activity

The cell bodies of primary sensory neurons that convey afferent signals from the periphery to the spinal dorsal horn reside in the DRGs (Figure 2A). Having established robust intrinsic circadian oscillations in the spinal cord dorsal horn, we next asked whether circadian clock activity is also present at the level of primary sensory neurons themselves. To address this, we cultured intact individual DRGs *ex vivo* and monitored PER2 reporter bioluminescence over multiple days under constant conditions. Circadian oscillations were readily detected in cervical, thoracic, and lumbar DRG explants (sacral ganglia were not assayed) (Figure 2B). Intrinsic circadian periods did not differ significantly across spinal levels (cervical: 24.67 ± 0.51 h; thoracic: 24.74 ± 0.24 h; lumbar: 24.52 ± 0.28 h; p > 0.05, Kruskal-Wallis test), indicating conserved circadian timing properties across DRGs.

**Figure 2.**
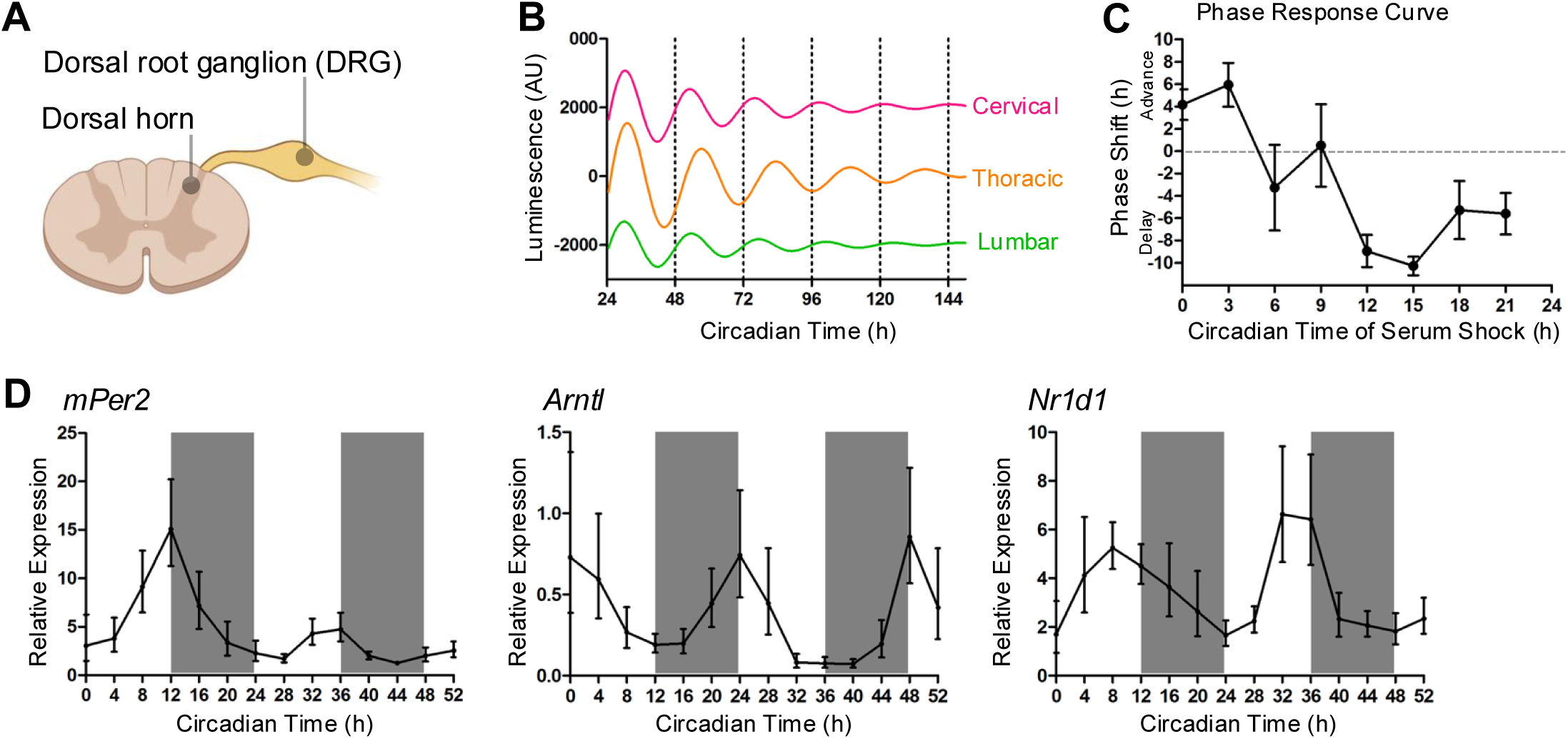
Autonomous circadian clock activity in the dorsal root ganglia. (A) Schematic of a transverse spinal cord section illustrating the dorsal horn and associated dorsal root ganglion (DRG). (B) Representative PER2 bioluminescence rhythms from individual mPer2Luc DRG explants at cervical, thoracic, and lumbar spinal levels under constant ex vivo conditions. (C) Phase–response curve for DRG explants following serum shock applied at different circadian times. Phase advances are plotted as positive values and phase delays as negative values. (D) Quantitative RT–PCR analysis of canonical circadian clock gene expression (mPer2, Arntl, and Nr1d1) in DRGs collected across a 52-hour time course under constant darkness. Data represent mean ± SEM.

A defining property of functional circadian clocks is their ability to reset phase in response to external stimuli in a phase-dependent manner. To assess whether DRG clocks exhibit this property, we applied a serum shock to DRG explants at different circadian times and quantified the resulting phase shifts. Phase resetting depended strongly on stimulus timing, yielding a characteristic phase-response relationship (Figure 2C). Serum shock administered early in the circadian cycle produced maximal phase advances (up to ∼6 h), whereas delivery at the opposite phase induced pronounced phase delays (up to ∼10 h), demonstrating that DRG clocks respond dynamically to resetting cues.

While *mPer2^Luc^* bioluminescence provides a sensitive readout of circadian clock activity *ex vivo*, we next sought to confirm that circadian rhythms are also present in DRGs *in vivo*. To this end, we collected DRGs across a 52-hour time course under constant darkness (DD), with sampling at 4-hour intervals (n = 3 animals per time point). Quantitative RT-PCR (qPCR) analysis revealed robust circadian expression of canonical clock genes *mPer2*, *Arntl* (*Bmal1*), and *Nr1d1* (Figure 2D). As expected, *mPer2* and *Arntl* transcripts oscillated in antiphase, consistent with established clock gene regulation ^35^. Together, these findings demonstrate that DRGs harbor self-sustained circadian clocks that operate autonomously *ex vivo* and are maintained *in vivo*.

### RNA-seq reveals phase-structured circadian gene expression in DRGs

Given the robust circadian rhythmicity in DRGs, we next asked whether circadian regulation extends broadly across the DRG transcriptome. To address this, we performed RNA sequencing on DRG samples collected across a 52-hour circadian time course under constant darkness (DD) conditions (CT0-CT52; n=3 mice per time point; Figure 3A). Following sequence alignment and normalization of FPKM values using the TuxedoTools pipeline ^26, 36^.

**Figure 3.**
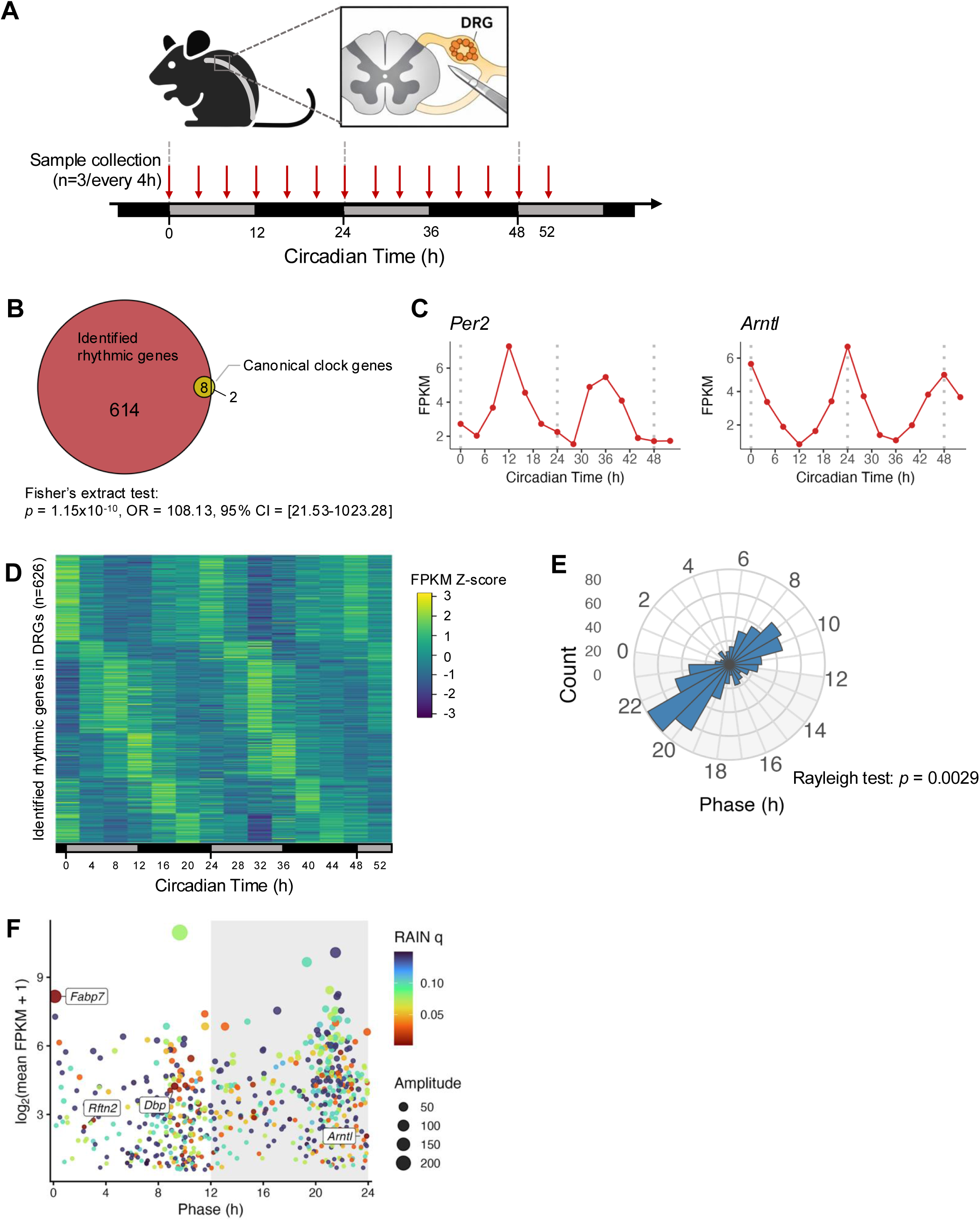
Comprehensive identification of circadian genes in dorsal root ganglia. (A) Schematic of the experimental design for circadian transcriptomic profiling of mouse dorsal root ganglia (DRGs). Adult mice were maintained under constant darkness (DD), and DRGs were collected every 4 h across a 52-h circadian time course (CT0–CT52; n = 3 mice per time point). The timeline indicates circadian time, with white and black bars denoting subjective day and night, respectively. DRG dissection from the spinal column is illustrated in the inset. (B) Enrichment of core clock genes among rhythmic DRG transcripts. Of the 626 circadian genes identified by RAIN (q < 0.15), 8 of 10 core clock genes (highlighted in yellow) were detected, significantly exceeding chance expectation (Fisher’s exact test: p = 1.15 × 10⁻¹⁰; OR = 108.13), indicating strong overrepresentation of clock components within the rhythmic gene set. (C) Representative expression profiles of Arntl and Per2 showing characteristic antiphase oscillations across circadian time. (D) Heatmap of z-scored expression profiles for RAIN-identified rhythmic genes in DRGs (n = 626), ordered by peak circadian phase to highlight temporal structure. (E) Circular histogram showing the distribution of peak circadian phases for rhythmic DRG genes identified by RAIN analysis (q < 0.15). Phase clustering was non-uniform as assessed by the Rayleigh test for circular uniformity (p < 0.01), indicating organized temporal regulation. Phases were derived from ARS-adjusted phase estimates and binned at 1-h resolution across the 24-h circadian cycle; the radial axis denotes gene count per phase bin. (F) Phase–expression plot of rhythmic DRG genes. Each point represents a rhythmic gene identified by RAIN (q < 0.15), plotted by peak circadian phase (x-axis) and mean transcript abundance [log₂(mean FPKM + 1); y-axis]. Point color indicates RAIN q-value, and point size represents oscillation amplitude. Gray shading denotes subjective night (CT12–24). Selected genes with strong rhythmicity are labeled.

To identify circadian transcripts in an assumption-free manner, we applied the RAIN algorithm to the time-series expression data ^27^. This analysis identified 626 genes exhibiting significant circadian rhythmicity (q < 0.15), representing 3.6% of the 17,394 expressed genes detected in DRGs (Supplementary File 1). To assess whether this number exceeded that expected by chance, we performed a permutation analysis in which circadian time labels were randomly reassigned ^30, 31^. The observed number of rhythmic genes (n = 626) lay far outside the null distribution generated from 1,000 permutations (empirical p = 0.000999; Supplementary Fig. 1A). This confirms that circadian rhythmicity in the DRG transcriptome reflects genuine temporal organization rather than statistical artifact. Canonical circadian clock genes were strongly enriched among the rhythmic transcripts (Fisher’s exact test: *p* = 1.15×10^−10^; OR = 108.13; 95% CI = 21.53-1023.28; Figure 3B). Eight of ten core clock genes, including *Arntl* (*Bmal1*) and *Per2*, exhibited robust rhythmic expression with their expected phase relationships (Figure 3C, Supplementary Figure 1B). These observations provide internal validation of both the experimental design and rhythmicity detection approach.

Phase-ordered visualization of rhythmic genes revealed clear temporal structure in DRG transcript oscillations. Hierarchical clustering of z-scored expression profiles grouped rhythmic transcripts according to their peak-phase preferences across the circadian cycle (Figure 3D). Consistent with this organization, peak-phase quantification revealed a non-uniform phase distribution: 262 genes (41.9%) peaked during the subjective day (CT0-CT12), whereas 364 genes (58.1%) peaked during the subjective night (CT12-CT24. This bimodal phase distribution was statistically non-random (Rayleigh test: p = 0.0029; Figure 3E), indicating that rhythmic gene expression in DRGs is organized into discrete, phase-locked transcriptional programs rather than being uniformly distributed across the circadian cycle.

To further characterize the relationship between rhythmic phase, expression level, and oscillation strength, we visualized individual rhythmic genes in phase–expression space (Figure 3F). Each gene was plotted according to its circadian phase and mean transcript abundance, with point color encoding RAIN significance (q-value) and point size representing oscillation amplitude. Rhythmic genes spanned a broad range of baseline expression levels and oscillation amplitudes, indicating that circadian regulation in DRGs is not confined to either highly abundant or low-expression transcripts, and is therefore unlikely to reflect stochastic noise or expression-level bias.

Clock-associated genes such as *Arntl* and *Dbp* exhibited peak phases consistent with established circadian timing. Among the most strongly rhythmic genes, *Fabp7* exhibited the lowest RAIN q-value, together with high oscillation amplitude and robust expression levels. *Fabp7*, which encodes fatty acid-binding protein 7, has been implicated in nervous system function and modulation of pain sensitivity ^37^, suggesting potential phase-dependent roles within the primary sensory axis. In addition, *Rftn2*, a gene with known sensory relevance ^38^, also exhibited robust oscillations with a preferential early daytime peak phase. Collectively, these findings are compatible with the idea that circadian timing in DRGs extends beyond core clock maintenance and organizes the transcriptome into phase-structured programs that include genes with established relevance to sensory biology.

### Circadian phase stratification reveals functional organization of rhythmic DRG genes

To investigate whether rhythmic transcripts are organized into phase-specific biological programs, we performed Gene Ontology (GO) analysis on circadian genes stratified by peak phase (Figure 4; Supplementary File 2). Phase binning was guided by circular variance analysis across multiple bin widths, and the distribution of circadian phases was visualized using kernel density estimation to highlight preferred phase windows (Figure 4A). Genes peaking during the early subjective day (CT0-3) exhibited enrichment for biological processes related to organic acid transport and carboxylic acid transport (Figure 4B, C), pathways relevant to neuronal metabolism and signaling ^39^. These enrichments were driven by genes including *Fabp7*, *Slc6a13*, and *Slc47a1*.

**Figure 4.**
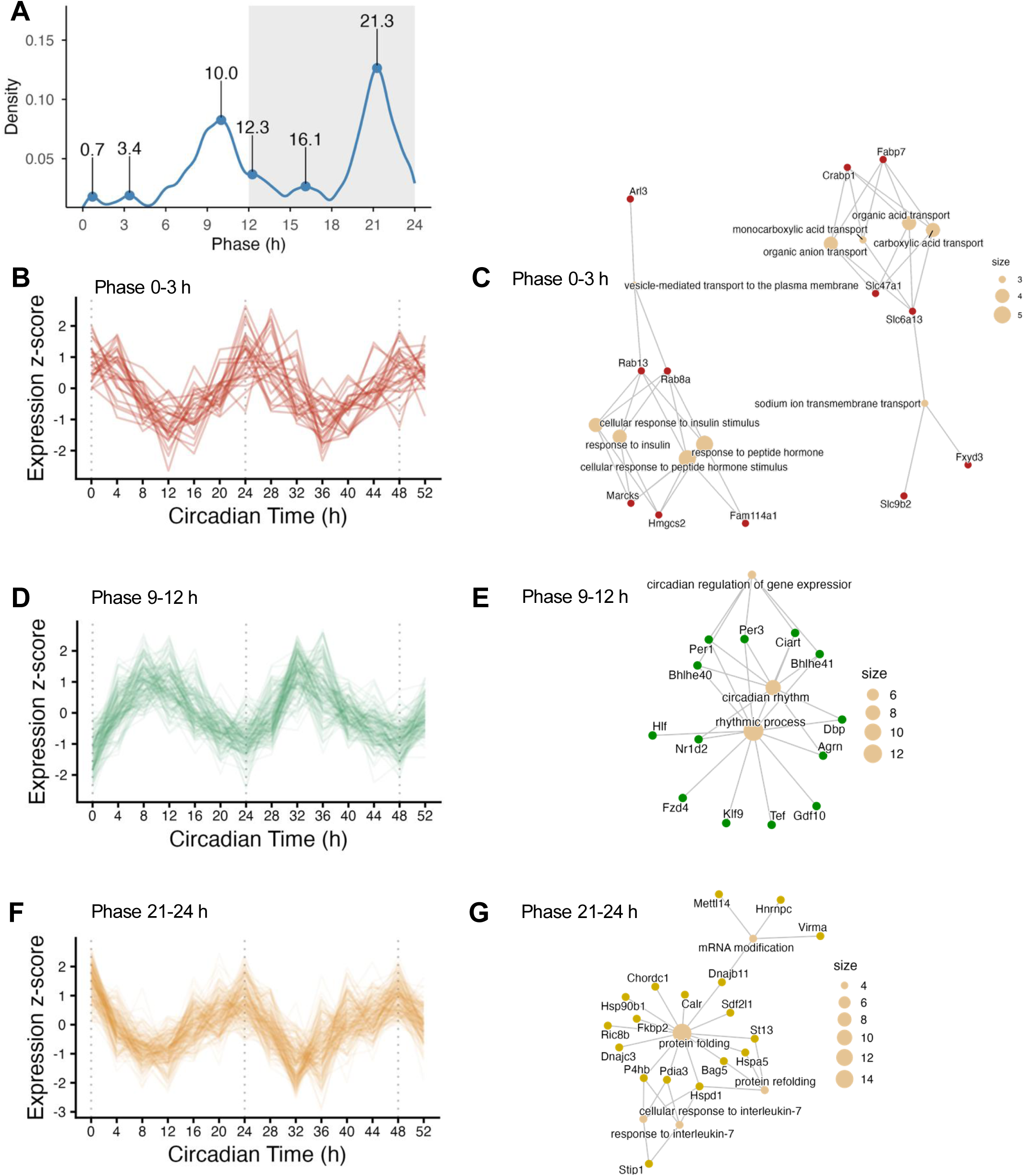
Circadian transcription in DRGs is organized into discrete phase-specific gene expression programs with distinct biological functions. (A) Kernel density estimate of peak circadian phases for rhythmic DRG transcripts identified by RAIN (q < 0.15). Local maxima indicate preferred phase windows, supporting stratification into discrete 3-h phase bins. (B, D, F) Z-score–normalized expression trajectories of rhythmic genes peaking at CT0–3 (B), CT9–12 (D), and CT21–24 (F). Each line represents an individual gene; dotted vertical lines denote 24-h intervals. (C, E, G) Gene Ontology biological process (GO-BP) enrichment networks for genes within the corresponding phase bins shown at left. Nodes represent enriched GO terms (beige) and associated genes (colored), with node size proportional to gene count per term. Terms meeting an exploratory enrichment threshold (BH-adjusted p < 0.1) are shown.

During mid-day phases (CT3-6 and CT6-9), GO analysis revealed enrichment of processes associated with neuronal structure and differentiation, including dendritic spine morphogenesis and neuron projection development (Supplementary Figure 2A-D), consistent with temporally restricted regulation of plasticity-related programs. In the late daytime transition (CT9-12), rhythmic genes were enriched for circadian regulation of gene expression and rhythmic processes themselves (Figure 4D, E), reflecting engagement of canonical clock-controlled transcriptional activity. In contrast, genes peaking during the subjective night exhibited less coherent functional annotation across phase bins. While early night phase (CT12-15) exhibited amino acid transport (Supplementary Figure 2E, F), no GO biological process terms reached statistical significance for genes peaking at CT15-18 or CT18-21 after multiple-testing correction. Notably, however, genes peaking in the late subjective night (CT21-24) were strongly enriched for protein folding, protein refolding, and cellular stress–response pathways (Figure 4F, G). This late-phase program indicates circadian coordination of proteostatic and RNA-regulatory processes.

Together, these biological network analyses indicate that circadian transcriptional programs in DRGs are organized into discrete, phase-specific functional programs rather than uniform oscillations across the circadian cycle. These findings suggest that circadian regulation in DRGs coordinates distinct biological functions at specific circadian phases, extending beyond core clock maintenance to shape temporal aspects of sensory and metabolic regulation.

### Baseline circadian organization reveals functional stratification of neuropathic pain-associated genes in DRGs

Given the robust time-dependent functional organization of the DRG transcriptome under baseline conditions, we next asked whether this temporal architecture reveals additional structure when examined across distinct biological contexts. Focusing on a condition directly relevant to somatosensory function, neuropathic pain, we cross-referenced baseline circadian DRG transcripts with gene sets associated with this condition.

Circadian genes identified by RAIN analysis (q < 0.15) were cross-referenced with previously reported gene sets associated with spared nerve injury (SNI)-induced neuropathic pain in DRGs ^40^ (Figure 5A, B) and neuropathic pain-related signaling pathways (PathCards version 5.26) annotated for spinal cord dorsal horn neurons (Figure 5C). Across both comparisons, only a small subset of neuropathic pain-associated genes exhibited significant circadian rhythmicity under baseline conditions. Enrichment analysis did not reveal statistically significant overrepresentation of circadian genes within neuropathic pain gene sets, indicating limited overlap between baseline circadian transcription and neuropathic pain-associated gene expression programs.

**Figure 5.**
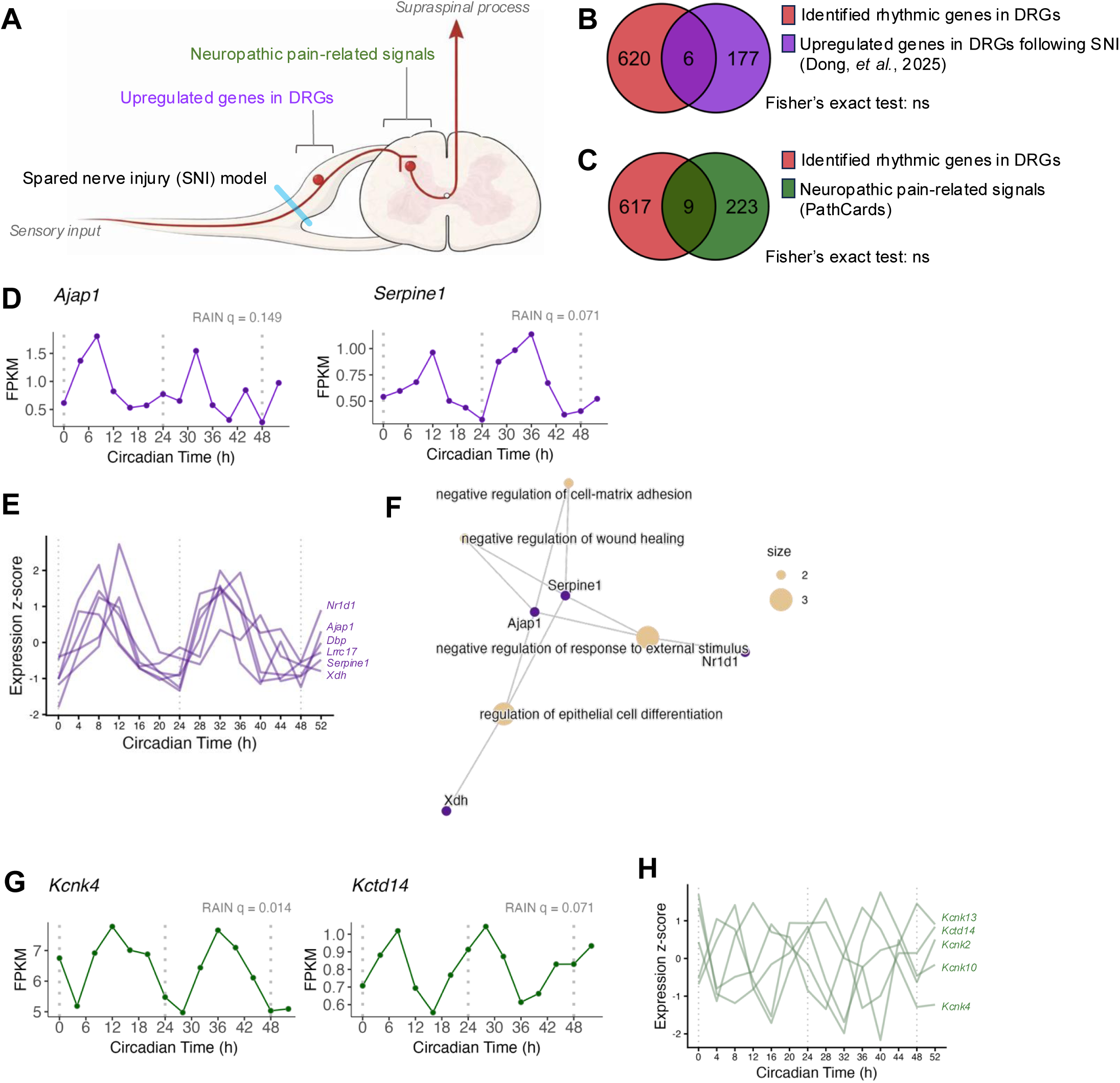
Baseline circadian organization reveals functional stratification of neuropathic pain–associated gene programs in DRGs. (A) Conceptual schematic of the spared nerve injury (SNI) model highlighting the sensory neuron–dorsal root ganglion (DRG)–spinal cord axis. Neuropathic pain–related signaling pathways (green) and genes reported to be upregulated in DRGs following nerve injury (purple) are shown in relation to ascending supraspinal processing, providing anatomical context for integration with baseline circadian transcriptional programs. (B) Venn diagram showing overlap between circadian rhythmic genes identified in DRGs under baseline conditions (RAIN q < 0.15; red) and genes upregulated in DRGs following neuropathic pain in published SNI datasets (purple; Dong et al., 2025). Only a small subset of genes overlapped (n = 6; Fisher’s exact test, not significant), indicating minimal enrichment of injury-induced genes among baseline circadian transcripts. (C) Venn diagram showing overlap between baseline circadian rhythmic DRG genes (RAIN q < 0.15; red) and neuropathic pain–related signaling genes curated from PathCards for dorsal horn neurons (green). As in panel B, overlap was limited (n = 9; Fisher’s exact test, not significant). (D) Representative baseline time-course expression profiles of overlapping rhythmic genes (Ajap1 and Serpine1) in DRGs under constant darkness. (E) Z-score–normalized baseline expression profiles of all overlapping rhythmic genes across the circadian cycle, illustrating convergence onto a shared circadian phase window. (F) Gene Ontology (GO) enrichment network for overlapping rhythmic genes, highlighting pathways related to tissue remodeling and regulation of injury responses. (p < 0.05, q < 0.05) (G) Representative baseline time-course expression profiles of neuropathic pain–associated potassium channel–related genes (Kcnk4 and Kctd14) identified from dorsal horn–centered pathway annotations. (H) Z-score–normalized baseline expression profiles of potassium channel–related genes (Kcnk2, Kcnk4, Kcnk10, Kcnk13, and Kctd14) across the circadian cycle, showing circadian modulation without clear phase convergence.

Genes shared between circadian and neuropathic pain-associated datasets included *Ajap1* and *Serpine1*, both of which have been implicated in regenerative or repair-associated processes following injury ^41^ (Figure 5B, D; Supplementary Figure 3A). Examination of their baseline expression profiles revealed preferential alignment to a shared circadian phase window (Figure 5E). As shown in earlier analyses, phase-convergent rhythmic genes in DRGs tend to participate in related biological functions rather than representing random assortments of oscillating transcripts. Consistent with this framework, Gene Ontology analysis of these overlapping genes highlighted pathways related to tissue remodeling and regulation of injury responses (Figure 5F), suggesting that these pathways are temporally organized under basal conditions, positioning them for coordinated engagement when recruited during neuropathic pain.

By contrast, when we cross-referenced neuropathic pain-associated signaling pathways in dorsal horn neurons, we identified multiple potassium channel-related transcripts (including *Kcnk4*, *Kcnk10*, *Kcnk13*, and *Kctd14*), which displayed circadian variation in DRGs under baseline conditions but did not show clear phase convergence (Figure 5C, G, H; Supplementary Figure 3B). The absence of phase clustering indicates that excitability-related components are temporally regulated in a gene-specific manner rather than organized into a single phase-restricted transcriptional program, despite belonging to the same functional family. This pattern is consistent with the functional diversity of potassium channel subtypes, which differ in their biophysical properties, regulatory mechanisms, and contributions to sensory neuron excitability ^42–44^. Such diversity suggests that distinct potassium channels may contribute to sensory processing at different circadian times. Collectively, these results indicate that neuropathic pain-associated genes map onto baseline circadian architecture in a function-dependent manner: remodeling-related genes align to shared phase windows, whereas excitability genes show subtype-specific timing without phase convergence.

## Discussion

Our findings indicate that the spinal dorsal horn and dorsal root ganglia harbor self-sustained circadian clocks, and that DRG gene expression is organized into phase-resolved transcriptional programs, supporting a model in which intrinsic clocks in the primary sensory axis structure daily windows of cellular preparedness. Functionally, this spans metabolic, transcriptional, and proteostatic states that bias how sensory tissues respond to sustained activity, inflammation, or injury.

Circadian function in the spinal cord and DRGs has remained incompletely characterized. Circadian rhythms and sleep in the spinal cord and DRGs have been implicated in pain ^12^, sensory dysesthesias such as restless legs syndrome ^45, 46^, and autoimmune disorders ^47^. However, much of the existing evidence comes from behavioral or anatomical studies conducted under light-dark cycles, making it difficult to disentangle intrinsic circadian regulation from behavioral or environmental influences. Here, by combining long-term *ex vivo* bioluminescence imaging with transcriptomic profiling under constant conditions, we provide direct evidence that both the spinal dorsal horn and DRGs harbor autonomous circadian clocks. The persistence of rhythmicity *ex vivo* demonstrates that these oscillations are tissue-intrinsic and not dependent on ongoing central or environmental cues. To our knowledge, this represents the first real-time characterization of PER2 rhythms in the spinal dorsal horn and DRGs, as well as a comprehensive circadian transcriptome of DRGs, and builds on prior work characterising these processes at discrete time-points in the context of chemotherapy-induced neuropathy ^21^.

Although expression of core clock genes has been reported previously in the spinal cord, the pronounced rhythmic amplitude observed in the dorsal horn and DRGs is notable. Within the spinal cord, dorsal horn regions, where primary afferent input is received and integrated, exhibited substantially higher circadian amplitude than ventral motor regions. Circadian oscillations have been described in spinal astrocytes, particularly at the lumbar levels, and have been linked to motor and autonomic regulation ^48, 49^. Our data extend these observations by showing that sensory-associated spinal compartments display especially robust circadian dynamics, emphasizing an underappreciated role for intrinsic clocks in shaping molecular programs within somatosensory pathways prior to extensive supraspinal processing.

A central advance of this study is the phase-structured organization of the DRG transcriptome. Although only 3.6% of expressed genes met our rhythmicity threshold, rhythmic genes were non-uniformly distributed across circadian phase and assembled into discrete temporal clusters. Importantly, GO enrichment within individual clusters revealed strong, coherent pathway structure. This pattern argues against a single global circadian output and instead supports multiple temporally separated transcriptional programs with distinct biological roles.

Genes peaking at early day (CT0-3) were enriched for organic acid and carboxylic acid transport, suggesting circadian regulation of metabolic substrate handling and signaling capacity. Several drivers of this enrichment have prior links to nervous system physiology. *Fabp7*, fatty acid–binding protein 7, is implicated in lipid trafficking and neuroinflammatory pathways, processes increasingly recognized as contributors to persistent pain states via immune-glial interactions and altered membrane signaling ^50^. A recent synthesis of FABP biology in pain highlights fatty-acid binding proteins as modulators of inflammatory and neuropathic pain-relevant processes ^51^, consistent with the idea that rhythmic *Fabp7* reflects time-of-day variation in lipid-handling capacity rather than acute sensory transmission. *Slc6a13* (GAT-3), a GABA transporter, provides a plausible mechanistic bridge between circadian transcription and longer-term excitability balance, because transporter expression influences ambient inhibitory tone and metabolite flux in neural tissue ^52^.

In mid-day (CT3-9), rhythmic genes are consistent with growth-factor and extracellular matrix (ECM)/regenerative biology, such as TGFβ/BMP-related signaling and ECM-associated genes such as *Igfbp5, Comp, Col6a1, Bmp6, Fbn1,* and *Fzd7* ^53, 54^. ECM remodeling and growth-factor signaling are well-established components of nerve injury responses and persistent pain ^55, 56^. Wnt signaling, in particular, has been shown to sensitize peripheral sensory neurons and contribute to pain hypersensitivity ^57^, supporting the relevance of a rhythmic Wnt-receptor axis including *Fzd* family members, in the primary afferent system. *Tiam1* ^58^*, Ngef* ^59^*, Il1rapl1* ^60^*, Nfatc4* ^61^, and *Abi3bp* ^62^ likely reflect shared molecular machinery involved in neurite remodeling, synapse-related scaffolding, or cytoskeletal regulation that can be deployed in multiple neuronal compartments. Notably, TIAM1-dependent plasticity has been mechanistically linked to chronic pain states, demonstrating that TIAM1-regulated structural programs can contribute to persistent pain phenotypes ^58^. Thus, rhythmic expression of structural/ECM and cytoskeletal regulators in DRGs may indicate circadian windows when remodeling capacity is transcriptionally favored. This idea that is particularly relevant for long-term circuit and axonal adaptations.

The most prominent night-associated enrichment emerged late in the cycle (CT21–24), with strong representation of protein folding and ER chaperone pathways such as *Hspa5/GRP78, Hsp90b1/GRP94, Calr, Pdia3, P4hb, Dnajc3*, and co-chaperones ^63, 64^. ER stress responses in DRGs have been directly implicated in neuropathic pain and mechanical hypersensitivity ^65^, including evidence that modulating ER-stress pathways in DRG neurons can alter pain behaviors ^66^. This phase bin also contained genes linked to mRNA modification/processing including *Mettl14, Virma/KIAA1429, Hnrnpc*, suggesting that circadian organization in DRGs extends beyond transcription into post-transcriptional control layers. METTL14-mediated m6A regulation has been identified as a contributor to chemotherapy-induced neuropathic pain ^67^, supporting the plausibility that rhythmic m6A machinery could affect longer-term transcript fate in sensory pathways. Together, these findings point to a late-cycle transcriptional emphasis on proteostasis and RNA regulation, potentially reflecting cumulative cellular demand across the activity cycle.

From a clinical translation perspective, these data provide a molecular scaffold for understanding why pain states and treatment responses can vary by time of day. Rather than implying that circadian clocks acutely gate pain perception, our results support a chronic pain-relevant model: circadian programs may bias the baseline physiological state of primary sensory neurons (metabolic substrate handling, growth-factor/ECM signaling, proteostasis, and RNA regulation), thereby influencing how the system responds to sustained inflammation, nerve injury, or repeated nociceptive input.

Overlapping genes, including *Ajap1*, *Serpine1*, *Lrrc17*, and *Xdh,* aligned with baseline phases enriched for transcriptional programs related to structural maintenance and remodeling. *Serpine1*, a regulator of plasminogen activator activity, has been reported to increase after nerve injury ^68^ and regulates plasmin-mediated extracellular matrix (ECM) proteolysis, a process implicated in axonal regeneration ^41^ and reduced when uPA/tPA are absent ^69^. Interestingly, the clock gene *Bmal1* (*Arntl*) has been reported to gate axon regeneration in DRG neurons, suggesting direct clock control of this process ^70^. *Ajap1* has been linked to Schwann cell myelination ^71^, leucine-rich repeat-containing proteins have been implicated in nociceptive circuit maintenance ^72–74^, and *Xdh* connects redox state to sensory function in DRGs ^75^. Together, these genes point to a pre-existing time window in which maintenance and remodeling-associated mechanisms are transcriptionally engaged under baseline conditions.

In addition, potassium channel-related components highlighted in dorsal horn neuropathic pain signaling pathways also showed baseline circadian modulation in DRGs. Potassium channels are key regulators of neuronal excitability and sensory signal integration in the dorsal horn, where peripheral input is relayed for central processing ^76^. Notably, unlike regenerative gene programs that converged on defined circadian phase windows, excitability-related transcripts did not collapse into a single phase-restricted pattern. Instead, individual potassium channel subtypes displayed gene-specific temporal profiles. In neuropathic pain models, this structured temporal modulation may be attenuated, with excitability-related genes showing more persistent activation, consistent with pathological engagement of sensory circuits ^77–79^.

Collectively, these findings support a model in which distinct molecular mechanisms within the primary sensory circuit are temporally organized into multiple circadian windows under normal conditions. Neuropathic pain models may accentuate specific subsets of these programs, but the underlying temporal architecture likely reflects broader physiological requirements of the sensory system. This framework highlights circadian phase as a determinant of the molecular state of DRGs and dorsal horn circuits and provides a rationale for exploring time-dependent vulnerability, resilience, and therapeutic responsiveness in sensory and pain-related disorders.

Several limitations should be noted. DRGs differ anatomically and functionally along the rostrocaudal axis, with cervical and lumbar ganglia predominantly innervating distal limbs and thoracic and sacral ganglia having greater autonomic involvement. Additionally, our transcriptome approach pooled DRGs across cell types at the cervical level, potentially obscuring level-specific or cell-type–specific rhythms. In addition, rhythmic mRNA does not necessarily imply rhythmic protein ^80, 81^, and rapid changes in excitability are unlikely to be driven solely by transcriptional oscillations. Future work combining single-cell/spatial transcriptomics with proteomic or phosphoproteomic profiling, and testing phase-dependent responses in nerve injury or inflammatory pain models, will be essential to connect these phase programs to causal mechanisms and therapeutic timing. Temperature is a well-established entrainment cue for peripheral circadian clocks ^82^. While we did not directly interrogate the molecular mechanisms underlying thermal entrainment in DRGs, previous work in peripheral tissue have implicated Heat Shock Factor 1 (HSF1) ^82^, as well as cold-induced RNA-binding proteins ^83^, in temperature-driven clock resetting. Given that DRGs are located outside the blood-brain barrier and experience local thermal fluctuations, temperature-dependent entrainment may represent a physiologically relevant mechanism for phase alignment *in vivo*, independent of photic input.

## Acknowledgements

A.B.R. was funded by the Perelman School of Medicine, University of Pennsylvania, the Wellcome Trust (100333/Z/12/Z), the European Research Council (ERC Starting Grant No. 281348, MetaCLOCK), the EMBO Young Investigators Programme, and the Lister Institute of Preventative Medicine. This work was supported also by NIH DP1DK126167 and R01GM139211 (to A.B.R.).

## Author contributions

S.Y. performed the data analysis, synthesized findings, and wrote the manuscript.

L.W. designed the project, performed experiments, data analysis and wrote initial drafts of the manuscript.

U.K.V. performed experiments and data analysis.

A.B.R. designed the project and analyzed results, secured funding, synthesized findings, and wrote the paper with contributions from the other authors.

All authors have reviewed the final manuscript and agree on its interpretation.

---

**Supplementary Figure 1.**
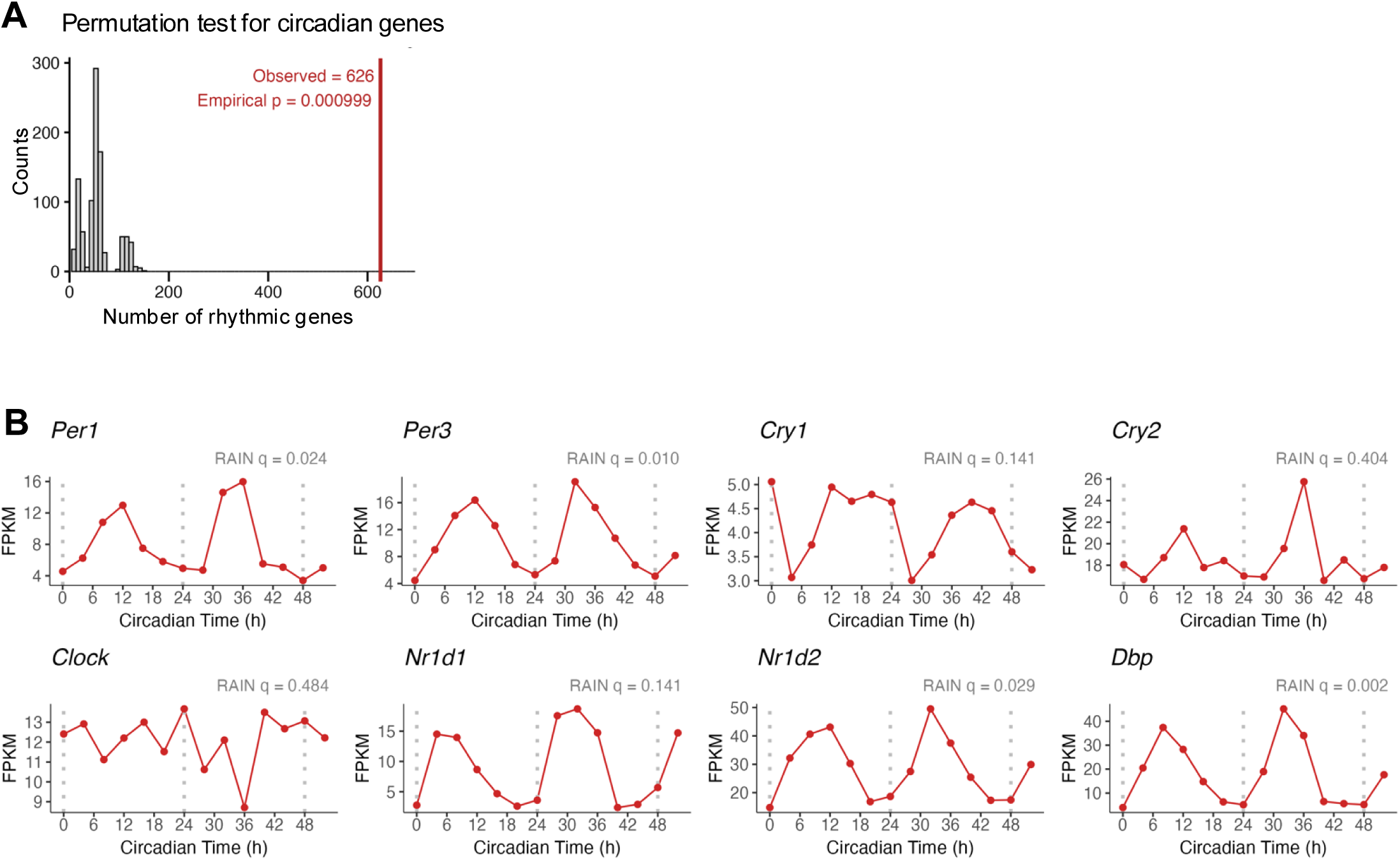
Validation of circadian rhythmicity in dorsal root ganglia. (A) Permutation-based significance testing of circadian rhythmicity in dorsal root ganglia (DRGs). Histogram shows the distribution of the number of rhythmic genes identified after random permutation of circadian time labels (1,000 permutations). The observed number of rhythmic genes in the real dataset (n = 626; red line) lies far outside the null distribution, yielding an empirical p-value of 0.000999 and indicating that rhythmic transcription in DRGs exceeds chance expectation. (B) Time-course RNA-seq expression profiles (FPKM) of canonical circadian clock genes (Per1, Per3, Cry1, Cry2, Clock, Nr1d1, Nr1d2, and Dbp) in mouse DRGs sampled across a 52-h circadian time course under constant darkness. Points represent mean expression at each time point, connected by lines for visualization. Vertical dashed lines denote successive circadian cycles. RAIN q-values are shown for each gene and reflect rhythmicity significance following multiple-testing correction. These profiles demonstrate heterogeneous rhythmic strength across individual clock components in DRGs. Supplementary Figure 2. Phase-resolved transcriptional programs in dorsal root ganglia.

**Supplementary Figure 2.**
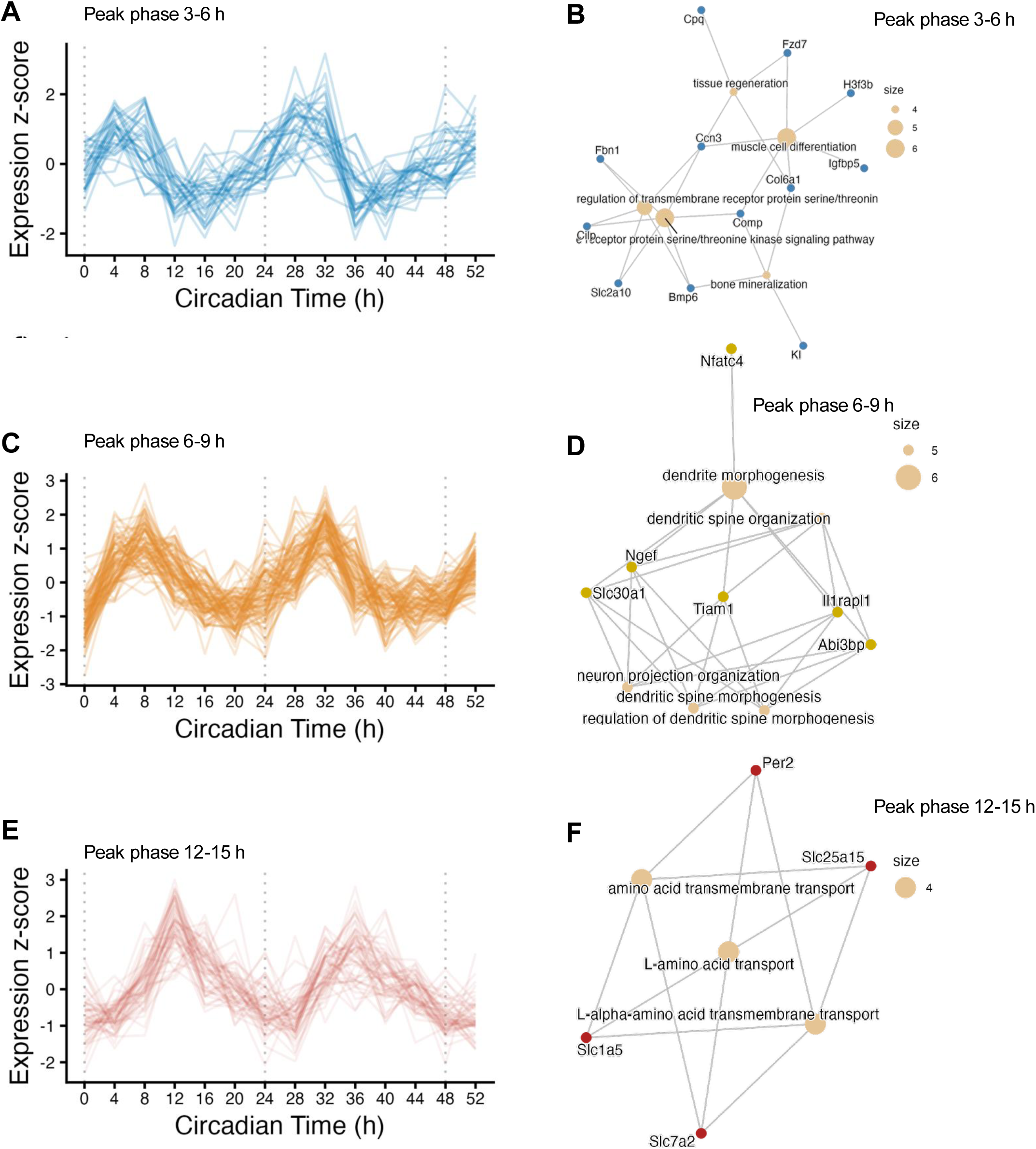
Phase-resolved transcriptional programs in dorsal root ganglia. (A, C, E) Z-score–normalized expression trajectories of rhythmic DRG genes binned by peak circadian phase, showing genes peaking at CT3–6 (A), CT6–9 (C), and CT12–15 (E). Each line represents an individual rhythmic gene identified by RAIN analysis (q < 0.15). Vertical dashed lines indicate circadian cycle boundaries (CT0, CT24, CT48). (B, D, F) Gene Ontology (GO) biological process enrichment networks corresponding to each phase bin shown at left. Nodes represent enriched GO terms (beige) and associated genes (colored), with node size proportional to the number of genes contributing to each term. Networks are shown for terms meeting an exploratory enrichment threshold (Benjamini–Hochberg–adjusted p < 0.1).

**Supplementary Figure 3.**
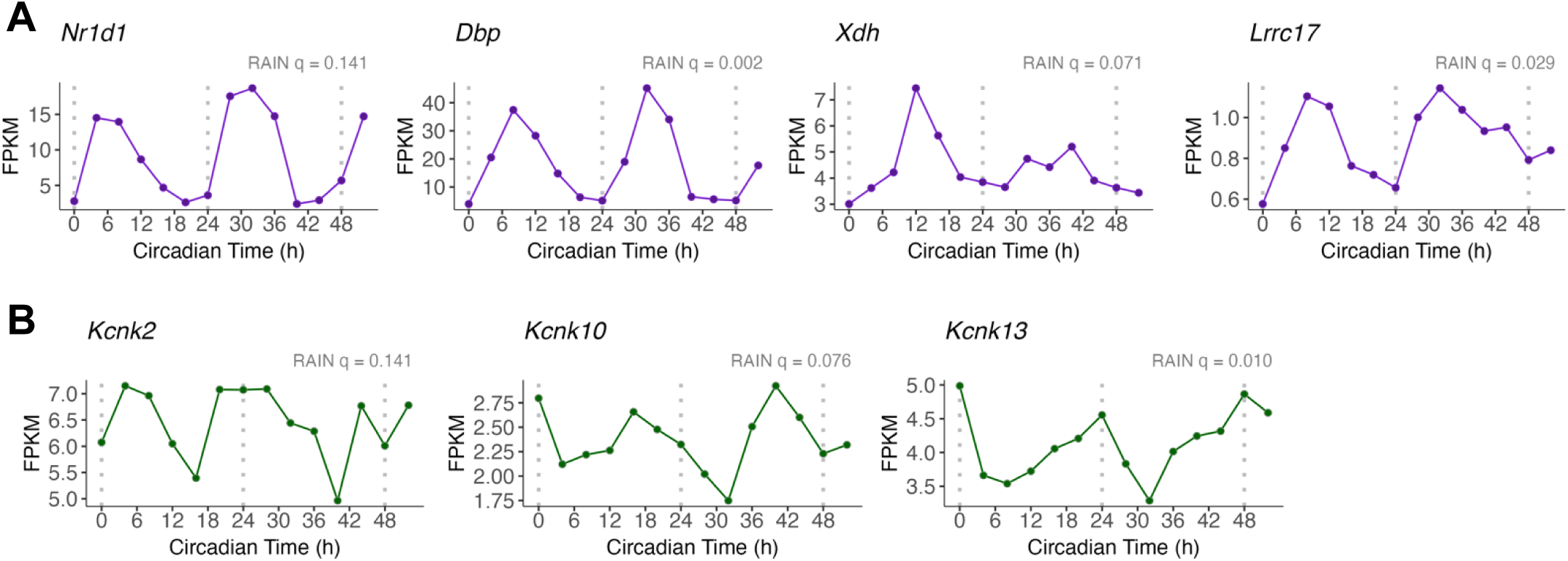
Baseline circadian modulation of neuropathic pain–associated genes in dorsal root ganglia. (A) Representative circadian expression profiles of genes overlapping between baseline circadian transcription and neuropathic pain–associated datasets (Nr1d1, Dbp, Xdh and Lrrc17) in DRGs under constant darkness. These genes exhibit defined circadian phase relationships under baseline conditions. RAIN q-values are shown for each gene. (B) Circadian expression profiles of potassium channel–related transcripts (Kcnk2, Kcnk10, and Kcnk13) implicated in neuropathic pain–associated signaling pathways. These genes display modest baseline circadian modulation in DRGs, consistent with temporal organization of excitability-related components under non-injured conditions.

**Supplementary File 1: Time-series DRG expression matrix and circadian rhythmicity metrics (RAIN/ARS).**

DRG RNA-seq time-course (CT0–CT52, 4-h resolution) with per-gene FPKM values and circadian rhythmicity metrics from RAIN and ARS, including phase/period estimates, amplitudes.

**Supplementary File 2: Phase-resolved Gene Ontology enrichment across circadian time in DRGs.**

Gene Ontology (GO) enrichment analyses for circadian transcripts in dorsal root ganglia (DRGs), stratified into consecutive 3-hour circadian time (CT) bins. Enrichment results are provided for Biological Process (BP), Molecular Function (MF), and Cellular Component (CC) categories.

## References

1. Reppert SM, Weaver DR (2002) Coordination of circadian timing in mammals. Nature 418:935–941

2. Hastings MH, Reddy AB, Maywood ES (2003) A clockwork web: circadian timing in brain and periphery, in health and disease. Nat Rev Neurosci 4:649–661

3. Jang K, Garraway SM (2024) A review of dorsal root ganglia and primary sensory neuron plasticity mediating inflammatory and chronic neuropathic pain. Neurobiol Pain 15:100151

4. Abraira VE, Ginty DD (2013) The Sensory Neurons of Touch. Neuron 79:618–639

5. Dong X, Dong X (2018) Peripheral and Central Mechanisms of Itch. Neuron 98:482–494

6. Lewin GR, Moshourab R (2004) Mechanosensation and pain. J Neurobiol 61:30–44

7. Ma Q (2012) Population coding of somatic sensations. Neurosci Bull 28:91–99

8. Friedman TN, Lamothe SM, Maguire AD, Hammond T, Tenorio G, Hilton BJ, Plemel JR, Kurata HT, Kerr BJ (2025) Plasticity of Mouse Dorsal Root Ganglion Neurons by Innate Immune Activation Is Influenced by Electrophysiological Activity. J Neurochem 169:e16292

9. Junker U, Wirz S (2010) Review Article: Chronobiology: influence of circadian rhythms on the therapy of severe pain. J Oncol Pharm Pr 16:81–87

10. Gilron I, Bailey JM, Vandenkerkhof EG (2013) Chronobiological Characteristics of Neuropathic Pain. Clin J Pain 29:755–759

11. Buttgereit F, Smolen JS, Coogan AN, Cajochen C (2015) Clocking in: chronobiology in rheumatoid arthritis. Nat Rev Rheumatol 11:349–356

12. Bruguerolle B, Labrecque G (2007) Rhythmic pattern in pain and their chronotherapy. Adv Drug Deliv Rev 59:883–895

13. Rasmussen NA, Farr LA (2003) Effects of Morphine and Time of Day on Pain and Beta-Endorphin. Biol Res Nurs 5:105–116

14. Wesche DL, Frederickson RCA (1979) Diurnal differences in opioid peptide levels correlated with nociceptive sensitivity. Life Sci 24:1861–1867

15. Crockett RS, Bornschein RL, Smith RP (1977) Diurnal variation in response to thermal stimulation: Mouse-hotplate test. Physiol Behav 18:193–196

16. Minett MS, Eijkelkamp N, Wood JN (2014) Significant Determinants of Mouse Pain Behaviour. Plos One 9:e104458

17. Chesler EJ, Wilson SG, Lariviere WR, Rodriguez-Zas SL, Mogil JS (2002) Influences of laboratory environment on behavior. Nat Neurosci 5:1101–1102

18. Kavaliers M, Hirst M, Teskey GC (1984) Aging and daily rhythms of analgesia in mice: Effects of natural illumination and twilight. Neurobiol Aging 5:111–114

19. Kavaliers M, Hirst M (1983) Daily rhythms of analgesia in mice: effects of age and photoperiod. Brain Res 279:387–393

20. Kusunose N, Koyanagi S, Hamamura K, Matsunaga N, Yoshida M, Uchida T, Tsuda M, Inoue K, Ohdo S (2010) Molecular Basis for the Dosing Time-Dependency of Anti-Allodynic Effects of Gabapentin in a Mouse Model of Neuropathic Pain. Mol Pain 6:1744–8069-6–83

21. Kim HK, Lee S-Y, Koike N, et al (2020) Circadian regulation of chemotherapy-induced peripheral neuropathic pain and the underlying transcriptomic landscape. Sci Rep 10:13844

22. Xia T, Cui Y, Qian Y, Chu S, Song J, Gu X, Ma Z (2016) Regulation of the NR2B-CREB-CRTC1 Signaling Pathway Contributes to Circadian Pain in Murine Model of Chronic Constriction Injury. Anesthesia Analg 122:542–552

23. Zhang J, Li H, Teng H, Zhang T, Luo Y, Zhao M, Li Y-Q, Sun ZS (2012) Regulation of Peripheral Clock to Oscillation of Substance P Contributes to Circadian Inflammatory Pain. Anesthesiology 117:149–160

24. Yoo S-H, Yamazaki S, Lowrey PL, et al (2004) PERIOD2::LUCIFERASE real-time reporting of circadian dynamics reveals persistent circadian oscillations in mouse peripheral tissues. Proc Natl Acad Sci 101:5339–5346

25. Hirota T, Lewis WG, Liu AC, Lee JW, Schultz PG, Kay SA (2008) A chemical biology approach reveals period shortening of the mammalian circadian clock by specific inhibition of GSK-3β. Proc National Acad Sci 105:20746–20751

26. Kim D, Pertea G, Trapnell C, Pimentel H, Kelley R, Salzberg SL (2013) TopHat2: accurate alignment of transcriptomes in the presence of insertions, deletions and gene fusions. Genome Biol 14:R36

27. Thaben PF, Westermark PO (2014) Detecting Rhythms in Time Series with RAIN. J Biol Rhythm 29:391–400

28. Yang R, Su Z (2010) Analyzing circadian expression data by harmonic regression based on autoregressive spectral estimation. Bioinformatics 26:i168–i174

29. Laloum D, Robinson-Rechavi M (2020) Methods detecting rhythmic gene expression are biologically relevant only for strong signal. Plos Comput Biol 16:e1007666

30. Sohn I, Owzar K, George SL, Kim S, Jung S-H (2009) A permutation-based multiple testing method for time-course microarray experiments. BMC Bioinform 10:336

31. Storey JD, Xiao W, Leek JT, Tompkins RG, Davis RW (2005) Significance analysis of time course microarray experiments. Proc Natl Acad Sci 102:12837–12842

32. Abe M, Herzog ED, Yamazaki S, Straume M, Tei H, Sakaki Y, Menaker M, Block GD (2002) Circadian Rhythms in Isolated Brain Regions. J Neurosci 22:350–356

33. Liu AC, Welsh DK, Ko CH, et al (2007) Intercellular Coupling Confers Robustness against Mutations in the SCN Circadian Clock Network. Cell 129:605–616

34. Osseward PJ, Pfaff SL (2019) Cell type and circuit modules in the spinal cord. Curr Opin Neurobiol 56:175–184

35. Oishi K, Sakamoto K, Okada T, Nagase T, Ishida N (1998) Antiphase Circadian Expression betweenBMAL1andperiodHomologue mRNA in the Suprachiasmatic Nucleus and Peripheral Tissues of Rats. Biochem Biophys Res Commun 253:199–203

36. Trapnell C, Roberts A, Goff L, Pertea G, Kim D, Kelley DR, Pimentel H, Salzberg SL, Rinn JL, Pachter L (2012) Differential gene and transcript expression analysis of RNA-seq experiments with TopHat and Cufflinks. Nat Protoc 7:562–578

37. Chen Y, Liu J, He Y, Lü Y, Yu W (2025) The Role of Fatty Acid Binding Protein 7 in Neurological Diseases. Mol Neurobiol 62:14801–14810

38. Faust TE, Gunner G, Schafer DP (2021) Mechanisms governing activity-dependent synaptic pruning in the developing mammalian CNS. Nat Rev Neurosci 22:657–673

39. Wang J, Geng B, Shen H-L, Xu X, Wang H, Wang C-F, Ma J-L, Wang Z-P (2012) Amino acid transport system A is involved in inflammatory nociception in rats. Brain Res 1449:38–45

40. Dong F-L, Yu L, Feng P-D, et al (2025) An atlas of neuropathic pain-associated molecular pathological characteristics in the mouse spinal cord. Commun Biol 8:70

41. Siconolfi LB, Seeds NW (2001) Induction of the plasminogen activator system accompanies peripheral nerve regeneration after sciatic nerve crush. J Neurosci : Off J Soc Neurosci 21:4336–47

42. Enyedi P, Czirják G (2010) Molecular Background of Leak K+ Currents: Two-Pore Domain Potassium Channels. Physiol Rev 90:559–605

43. Honoré E (2007) The neuronal background K2P channels: focus on TREK1. Nat Rev Neurosci 8:251–261

44. Natale AM, Deal PE, Minor DL (2021) Structural Insights into the Mechanisms and Pharmacology of K2P Potassium Channels. J Mol Biol 433:166995

45. Clemens S, Sawchuk MA, Hochman S (2005) Reversal of the circadian expression of tyrosine-hydroxylase but not nitric oxide synthase levels in the spinal cord of dopamine D3 receptor knockout mice. Neuroscience 133:353–357

46. Yamuy J, Fung SJ, Xi M, Chase MH (2010) State-dependent control of lumbar motoneurons by the hypocretinergic system. Exp Neurol 221:335–345

47. He J, Hsuchou H, He Y, Kastin AJ, Mishra PK, Fang J, Pan W (2014) Leukocyte infiltration across the blood-spinal cord barrier is modulated by sleep fragmentation in mice with experimental autoimmune encephalomyelitis. Fluids Barriers CNS 11:27

48. Morioka N, Sugimoto T, Tokuhara M, Nakamura Y, Abe H, Hisaoka K, Dohi T, Nakata Y (2012) Spinal astrocytes contribute to the circadian oscillation of glutamine synthase, cyclooxygenase-1 and clock genes in the lumbar spinal cord of mice. Neurochem Int 60:817–826

49. Sugimoto T, Morioka N, Sato K, Hisaoka K, Nakata Y (2011) Noradrenergic regulation of period1 expression in spinal astrocytes is involved in protein kinase A, c-Jun N-terminal kinase and extracellular signal-regulated kinase activation mediated by α1- and β2-adrenoceptors. Neuroscience 185:1–13

50. Wu C, Lin J, Chen Q, Zhao W, Kawahata I, Cheng A (2025) Targeting the FABP Axis: Interplay Between Lipid Metabolism, Neuroinflammation, and Neurodegeneration. Cells 14:1502

51. Bogdan DM, Studholme K, DiBua A, Gordon C, Kanjiya MP, Yu M, Puopolo M, Kaczocha M (2022) FABP5 deletion in nociceptors augments endocannabinoid signaling and suppresses TRPV1 sensitization and inflammatory pain. Sci Rep 12:9241

52. Orefice LL, Mosko JR, Morency DT, et al (2019) Targeting Peripheral Somatosensory Neurons to Improve Tactile-Related Phenotypes in ASD Models. Cell 178:867–886.e24

53. Chen G, Deng C, Li Y-P (2012) TGF-β and BMP Signaling in Osteoblast Differentiation and Bone Formation. Int J Biol Sci 8:272–288

54. Liao J, Wu T, Zhang Q, et al (2026) TGF-β/BMP signaling in skeletal biology: molecular mechanisms, regulatory networks, and therapeutic implications in development, regeneration, and disease. Bone Res 14:6

55. Metafune M, Muratori L, Fregnan F, Ronchi G, Raimondo S (2025) The extracellular matrix in peripheral nerve repair and regeneration: a narrative review of its role and therapeutic potential. Front Neuroanat 19:1628081

56. Gantus MAV, Nasciutti LE, Cruz CM, Persechini PM, Martinez AMB (2006) Modulation of extracellular matrix components by metalloproteinases and their tissue inhibitors during degeneration and regeneration of rat sural nerve. Brain Res 1122:36–46

57. Simonetti M, Agarwal N, Stösser S, et al (2014) Wnt-Fzd Signaling Sensitizes Peripheral Sensory Neurons via Distinct Noncanonical Pathways. Neuron 83:104–121

58. Li L, Ru Q, Lu Y, Fang X, Chen G, Saifullah AB, Yao C, Tolias KF (2023) Tiam1 coordinates synaptic structural and functional plasticity underpinning the pathophysiology of neuropathic pain. Neuron 111:2038–2050.e6

59. Lin W, Szaro BG (1996) Effects of Intermediate Filament Disruption on the Early Development of the Peripheral Nervous System ofXenopus laevis. Dev Biol 179:197–211

60. Gambino F, Pavlowsky A, Béglé A, et al (2007) IL1-receptor accessory protein-like 1 (IL1RAPL1), a protein involved in cognitive functions, regulates N-type Ca2+-channel and neurite elongation. Proc Natl Acad Sci 104:9063–9068

61. Li L, Ke K, Tan X, et al (2013) Up-regulation of NFATc4 Involves in Neuronal Apoptosis Following Intracerebral Hemorrhage. Cell Mol Neurobiol 33:893–905

62. Aiken J, Holzbaur ELF (2021) Cytoskeletal regulation guides neuronal trafficking to effectively supply the synapse. Curr Biol 31:R633–R650

63. McLaughlin M, Vandenbroeck K (2011) The endoplasmic reticulum protein folding factory and its chaperones: new targets for drug discovery? Br J Pharmacol 162:328–345

64. Petrova K, Oyadomari S, Hendershot LM, Ron D (2008) Regulated association of misfolded endoplasmic reticulum lumenal proteins with P58/DNAJc3. EMBO J 27:2862–2872

65. Inceoglu B, Bettaieb A, Silva CAT da, Lee KSS, Haj FG, Hammock BD (2015) Endoplasmic reticulum stress in the peripheral nervous system is a significant driver of neuropathic pain. Proc Natl Acad Sci 112:9082–9087

66. Zhang E, Yi M-H, Shin N, et al (2015) Endoplasmic reticulum stress impairment in the spinal dorsal horn of a neuropathic pain model. Sci Rep 5:11555

67. Lu W, Yang X, Zhong W, et al (2024) METTL14-mediated m6A epitranscriptomic modification contributes to chemotherapy-induced neuropathic pain by stabilizing GluN2A expression via IGF2BP2. J Clin Investig 134:e174847

68. Nilsson A, Moller K, Dahlin L, Lundborg G, Kanje M (2005) Early changes in gene expression in the dorsal root ganglia after transection of the sciatic nerve; effects of amphiregulin and PAI-1 on regeneration. Mol Brain Res 136:65–74

69. Siconolfi LB, Seeds NW (2001) Mice lacking tPA, uPA, or plasminogen genes showed delayed functional recovery after sciatic nerve crush. J Neurosci : Off J Soc Neurosci 21:4348–55

70. Halawani D, Wang Y, Ramakrishnan A, Estill M, He X, Shen L, Friedel RH, Zou H (2023) Circadian clock regulator Bmal1 gates axon regeneration via Tet3 epigenetics in mouse sensory neurons. Nat Commun 14:5165

71. Basak S, Desai DJ, Rho EH, Ramos R, Maurel P, Kim HA (2015) E-Cadherin enhances neuregulin signaling and promotes Schwann cell myelination. Glia 63:1522–1536

72. Schroeder A, Wit J de (2018) Leucine-rich repeat-containing synaptic adhesion molecules as organizers of synaptic specificity and diversity. Exp Mol Med 50:1–9

73. Bando T, Morikawa Y, Hisaoka T, Komori T, Miyajima A, Senba E (2012) Expression pattern of leucine-rich repeat neuronal protein 4 in adult mouse dorsal root ganglia. Neurosci Lett 531:24–29

74. Deng M, Chen S-R, Zhou M-H, Zhang J, Huang Y, Chen H, Benavides F, Sah R, Pan H-L (2025) LRRC8A constitutively inhibits pain hypersensitivity in rodent models by restraining NMDA receptor activity at spinal cord synapses. Sci Transl Med 17:eadu4879

75. Hsieh C-P (2008) Redox modulation of A-type K+ currents in pain-sensing dorsal root ganglion neurons. Biochem Biophys Res Commun 370:445–449

76. Flauaus C, Engel P, Zhou F, Petersen J, Ruth P, Lukowski R, Schmidtko A, Lu R (2022) Slick Potassium Channels Control Pain and Itch in Distinct Populations of Sensory and Spinal Neurons in Mice. Anesthesiology 136:802–822

77. Costigan M, Scholz J, Woolf CJ (2009) Neuropathic Pain: A Maladaptive Response of the Nervous System to Damage. Annu Rev Neurosci 32:1–32

78. Campbell JN, Meyer RA (2006) Mechanisms of Neuropathic Pain. Neuron 52:77–92

79. Sukumaran S, Almon RR, DuBois DC, Jusko WJ (2010) Circadian rhythms in gene expression: Relationship to physiology, disease, drug disposition and drug action. Adv Drug Deliv Rev 62:904–917

80. Robles MS, Cox J, Mann M (2014) In-Vivo Quantitative Proteomics Reveals a Key Contribution of Post-Transcriptional Mechanisms to the Circadian Regulation of Liver Metabolism. PLoS Genet 10:e1004047

81. Reddy AB, Karp NA, Maywood ES, et al (2006) Circadian Orchestration of the Hepatic Proteome. Curr Biol 16:1107–1115

82. Buhr ED, Yoo S-H, Takahashi JS (2010) Temperature as a Universal Resetting Cue for Mammalian Circadian Oscillators. Science 330:379–385

83. Schibler U (2009) The 2008 Pittendrigh/Aschoff Lecture: Peripheral Phase Coordination in the Mammalian Circadian Timing System. J Biol Rhythm 24:3–15

